# Chromatin changes in PIF-regulated genes parallel their rapid transcriptional response to light

**DOI:** 10.1101/2021.10.04.463089

**Authors:** Eduardo González-Grandío, Simón Álamos, Yu Zhang, Jutta Dalton-Roesler, Krishna K. Niyogi, Hernán G. García, Peter H. Quail

## Abstract

As sessile organisms, plants must adapt to a changing environment, sensing variations in resource availability and modifying their development in response. Light is one of the most important resources for plants, and its perception by sensory photoreceptors (e.g. phytochromes) and subsequent transduction into long-term transcriptional reprogramming have been well characterized. Chromatin changes have been shown to be involved in photomorphogenesis. However, the initial short-term transcriptional changes produced by light and what factors enable these rapid changes are not well studied. Here, we identify rapidly light-responsive, PIF (Phytochrome Interacting Factor) direct-target genes (LRP-DTGs). We found that a majority of these genes also show rapid changes in Histone 3 Lysine-9 acetylation (H3K9ac) in response to the light signal. Detailed time-course analysis of transcriptional and chromatin changes showed that, for light-repressed genes, H3K9 deacetylation parallels light-triggered transcriptional repression, while for light-induced genes, H3K9 acetylation appeared to somewhat precede light-activated transcription. However, real-time imaging of transcription elongation revealed that, in fact, H3K9 acetylation also parallels transcriptional induction. Collectively, the data raise the possibility that light-induced transcriptional and chromatin-remodeling processes are mechanistically intertwined. Histone modifying proteins involved in long term light responses do not seem to have a role in this fast response, indicating that different factors might act at different stages of the light response. This work not only advances our understanding of plant responses to light, but also unveils a system in which rapid chromatin changes in reaction to an external signal can be studied under natural conditions.

## Introduction

One of the most drastic changes during plant development is deetiolation, the switch from skotomorphogenesis (development in the dark) into photomorphogenesis (development in the light). This change not only implies switching from heterotrophy to autotrophy, but also includes several developmental changes such as reduced hypocotyl elongation, opening of the apical hook and greening of cotyledons (Schafer and Nagy, 2006; Franklin and Quail, 2010). Light signals that trigger deetiolation are perceived by photoreceptors. In Arabidopsis, phytochrome A (phyA) and phyB are the main sensors that regulate early photomorphogenesis (Franklin and Quail, 2010). Upon photoactivation, the active phy is translocated from the cytoplasm into the nucleus where it physically interacts with Phytochrome Interacting Factors (PIFs), inducing their rapid transphosphorylation, ubiquitination and proteasome-mediated degradation of the phy-PIF complex (Bauer et al., 2004; Nagatani, 2004; Al-Sady et al., 2006; Jiao et al., 2007; Bae and Choi, 2008; Pfeiffer et al., 2012; Ni et al., 2013, 2014, 2017). PIFs are a subfamily of bHLH transcription factors, comprising eight members in *Arabidopsis thaliana*. PIF1, PIF3, PIF4 and PIF5 (known as the “PIF quartet”) have a central role in maintaining the transcriptional program that underlies skotomorphogenic development (Leivar et al., 2008; Shin et al., 2009). The quadruple mutant for these four PIFs (*pifq*) displays a phenotype in total darkness that strongly resembles that of normal light-grown wild-type seedlings (Leivar et al., 2008; Shin et al., 2009).

To identify PIF-direct target genes (PIF-DTGs), our group analyzed the genome-wide binding profile of each of the PIF quartet members by Chromatin Immunoprecipitation-sequencing (ChIP-seq), and the corresponding transcriptomic profile of dark-grown wild type, *pifq*, single *pif* and triple *pif* mutants, by RNA sequencing (RNA-seq) (Hornitschek et al., 2012; Zhang et al., 2013; Pfeiffer et al., 2014). Integration of both datasets allowed the identification of 338 PIF-DTGs, genes whose promoter region is bound by one or more PIF quartet members at a G-box or PIF Binding Element (PBE), and whose transcript levels are misregulated in *pifq* mutant plants grown in the dark (Pfeiffer et al., 2014). General transcriptional reprogramming that results in photomorphogenesis, occurs upon light exposure of dark-grown plants, as a consequence of PIF degradation (Leivar et al., 2009). However, the initial dynamics of light-induced transcriptional changes of PIF-DTGs has not been studied.

In eukaryotes, chromatin structure modification is a key factor of transcription regulation (Venkatesh and Workman, 2015). Among the many histone post-translational modifications, acetylation seems to play a major role in this process (Jiang et al., 2020). Histone acetylation, catalyzed by histone acetyltransferases (HATs), has been associated with transcriptional activation, while histone deacetylation by histone deacetylases (HDACs) is linked to transcriptional repression (Pandey et al., 2002). Whether the role of histone modifications in transcriptional regulation is causal or consequential is not well understood (Henikoff and Shilatifard, 2011; Morgan and Shilatifard, 2020). Previous research has established that histone acetylation plays an essential role during photomorphogenesis (Barneche et al., 2014). Transcriptional regulation and development of light-grown plants is greatly altered in Histone 3 Lysine 9 acetylation (H3K9ac) deposition (HAG1/GCN5 and HAF2/TAF1) and removal mutants (HDA19/HD1) (Bertrand et al., 2005; Benhamed et al., 2006; Guo et al., 2008). Recent evidence suggests that PIF-exerted transcriptional regulation might be executed by altering the histone modification landscape. It has been described that PIF1 and PIF3 directly interact with HDA15 in the dark to repress the expression of germination and chlorophyl biosynthesis genes, respectively (Liu et al., 2013; Gu et al., 2017). PIF7 has also been shown to induce H3K9ac deposition on its downstream genes in response to changes in light quality (Peng et al., 2018; Martínez-García and Moreno-Romero, 2020; Willige et al., 2021). HY5, a key photomorphogenesis transcription factor also interacts with HDA15 and decreases histone acetylation of genes involved in hypocotyl elongation during photomorphogenesis to repress their expression (Zhao et al., 2019). Additionally, phyB has been shown to redundantly control chromatin remodeling to inhibit the transcriptional activation of growth-promoting genes by PIFs (Kim et al., 2021). It is still unclear if these factors that control histone acetylation in dark or light-grown plants are also involved in the very initial steps after light exposure that will trigger the photomorphogenesis transcriptional program.

Here, to explore the potential role of chromatin remodeling as an intermediary in light-triggered regulation of PIF-DTGs, we focused on H3K9 acetylation as a mark of transcriptionally active genes. We first identified rapid light-responding PIF-DTGs by comparing the transcriptomic profile of dark-grown seedlings with those of dark-grown *pifq* mutants and dark-grown seedlings after a short red-light treatment. We also profiled genome-wide H3K9ac localization in these plants. We found that the majority of light-responsive PIF-DTGs also showed changes in H3K9ac. We then conducted a detailed time-course analysis of mRNA and H3K9ac levels on selected light-responsive PIF-DTGs. This analysis initially suggested that the relationship between H3K9ac and transcription differs in light-repressed and light-induced PIF-DTGs. Real-time transcription initiation imaging, however, suggests instead that H3K9ac also parallels transcriptional induction in response to light.

## Results and discussion

### Identification of rapidly responding, light regulated PIF-direct target genes

In our previous work, we defined PIF-DTGs as genes that are miss-regulated in *pifq* mutant plants grown in the dark and whose promoter is bound by any of the PIF quartet proteins (Pfeiffer et al., 2014). In order to identify genes that respond most directly to changes in PIF abundance, we focused on genes whose transcript levels change rapidly in wild-type seedlings after a short exposure to red light, when the PIFs have been almost completely degraded (Bauer et al., 2004; Shen et al., 2005, 2007; Lorrain et al., 2007). For this purpose, we measured genome-wide steady state mRNA levels by RNA-sequencing in “true dark”-grown, wild-type seedlings (D) and in “true dark”-grown seedlings treated for one hour with red light (R1h). We also measured mRNA levels in seedlings grown in continuous white light (WLc) as a reference expression profile of plants grown during an extended light regimen (Figure 1A).

**Figure 1.**
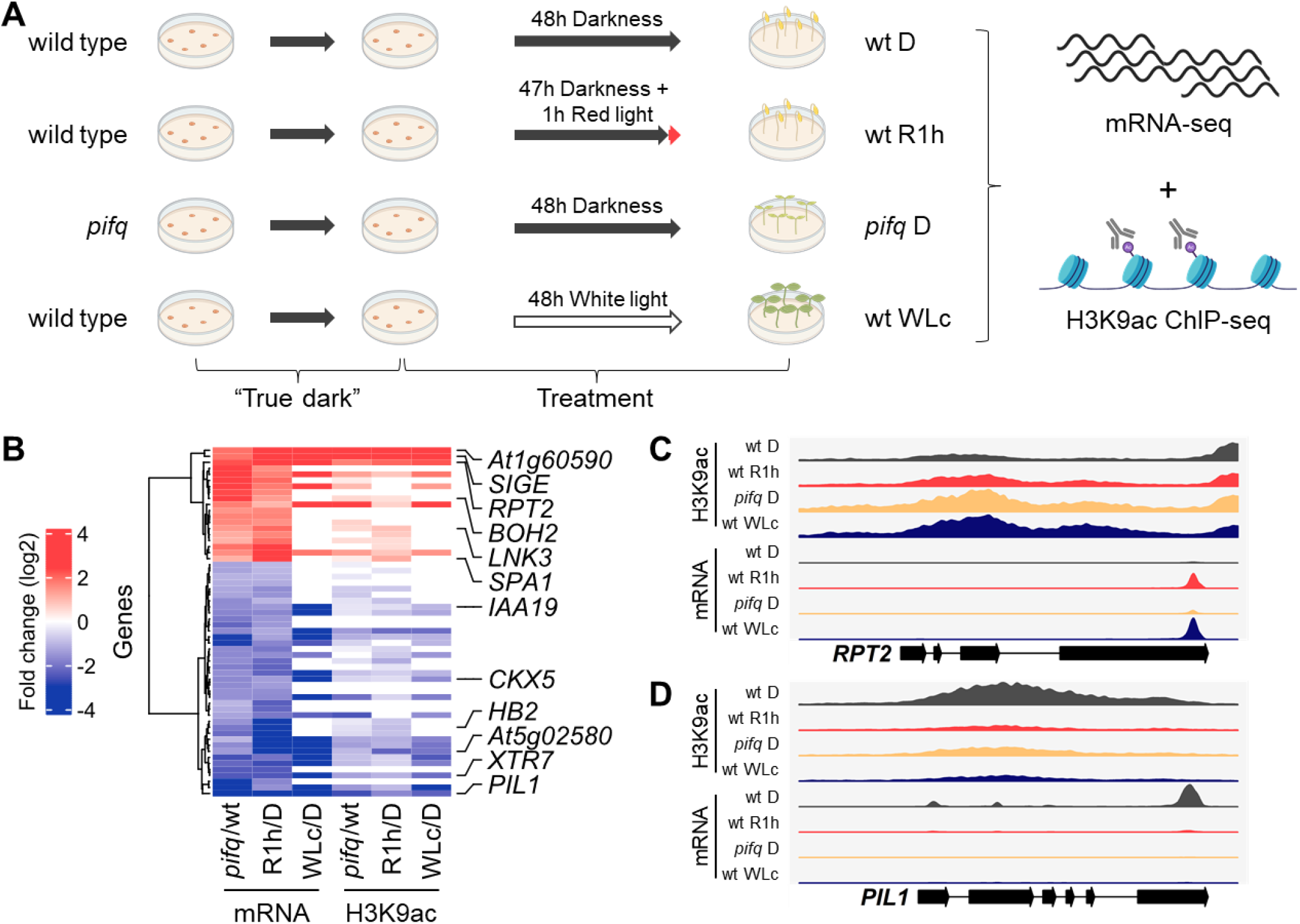
H3K9ac levels change in light-responsive PIF-DTGs (LRP-DTGs) both in *pifq* and in response to red light. A, Experimental setup. Wild type or *pifq* seedlings were grown in “true dark” conditions and treated for two days before sample collection and processing for mRNA- and H3K9ac ChIP-sequencing. B, Heat-map showing the changes in mRNA and H3K9ac levels comparing dark-grown *pifq* versus wild type, red-light treated (R1h) versus dark-grown wild type (D), and continuous white light-grown (WLc) vs dark grown wild-type seedlings. All the PIF-DTGs that showed statistically significant two-fold change in mRNA levels in the same direction in *pifq* vs wt and R1h vs D are shown. Six light-induced/PIF-repressed and six light-repressed/PIF-induced genes for further analyses are highlighted. C, Read mapping profile of H3K9ac ChIP-seq and RNA-seq in *RPT2*, a light-induced LRP-DTG. D, Read mapping profile of H3K9ac ChIP-seq and RNA-seq in *PIL1*, a light-repressed LRP-DTG. Note that RNA-seq was performed on 3’-end purified mRNA.

An initial analysis of R1h and WLc transcriptional profiles compared to D showed that the large transcriptional reprogramming occurring in WLc could be triggered by a small number of genes that change rapidly in response to the first exposure to light (R1h). These genes do not necessarily need to be continuously activated or repressed once the transcriptional reprograming has been initiated, our data showed that only somewhat over half of the genes that change initially in response to light exposure remain in that state after an extended light regime (Supplemental Dataset S2, Figure S1).

Surprisingly, after R1h, only 19% PIF-induced PIF-DTGs were repressed, and 15% of the PIF-repressed PIF-DTGs were induced (Figure S2). Previously identified PIF-DTGs that do not change rapidly in response to light could have a slower transcriptional response to PIF degradation, be indirectly regulated by them or could have been misidentified as PIF-DTGs due to non-functional PIF binding resulting in a lack of transcriptional regulation (see below). These possibilities are non-mutually exclusive and seem to be occurring. Using the WLc data, we could identify slower-responding PIF-regulated genes: 60% of PIF-induced PIF-DTGs are both R1h- and WLc-repressed, and 43% of PIF-repressed PIF-DTGs are R1h- and WLc-induced (Figure S3). Still, some PIF-DTGs do not show altered transcription either after R1h or WLc. Their altered transcriptional levels in *pifq* mutants grown in the dark could be caused by indirect effects (they could be downstream targets of PIF-DTGs). Additionally, these genes could represent cases where PIF-binding is non-functional and does not result in transcriptional regulation of the gene downstream of the PIF-binding site, as has been reported for other transcription factors in many ChIP-seq experiments (Biggin, 2011). These results reflect the limitation of using transcriptomic profiling of constitutive mutants in conjunction with genome wide binding assays to identify transcription factor direct target genes, leading to overestimation of the actual number of direct downstream genes. Complementing these studies with short-term response assays (in this case R1h) is essential to narrow down direct targets of transcription factors.

In summary, we have identified PIF-regulated genes whose expression changes very rapidly in response to light. These genes are directly regulated by PIFs and are likely candidates to effect the initial changes downstream of light perception that will trigger the photomorphogenesis developmental plan. For convenience, we will use the term “LRP-DTGs” (for <u>L</u>ight <u>R</u>esponsive <u>PIF</u> <u>D</u>irect <u>T</u>arget <u>G</u>enes) from here on to refer to the newly redefined genes that are bound by PIFs in the dark, have altered mRNA levels in the *pifq* mutants in the dark, and respond rapidly to R1h. Gene ontology enrichment analysis of light-repressed/PIF-induced LRP-DTGs revealed that they are enriched in transcription factors and in proteins known to be involved in red light, far-red light and auxin responses. These genes include *IAA19*, *IAA29*, *PIL1*, *PIL2*, and *HB2*. Light-induced/PIF-repressed LRP-DTGs were also enriched in proteins known to be involved in far-red, red and blue light responses, including genes such as *SPA1*, *RPT2*, *SIGE* and *LHCB2.4*.

### Chromatin changes shortly after red-light exposure

To investigate the possible involvement of rapid chromatin remodeling in the PIF-exerted regulation of LRP-DTGs, we conducted genome-wide profiling of H3K9ac by ChIP-seq in dark-grown wild type and *pifq* seedlings, and R1h-treated wild type seedlings. We also profiled H3K9ac in wild type seedlings grown in WLc for comparison as we did for transcriptome profiling (Figure 1A). We chose H3K9ac because it is a histone modification that has been shown to be associated with transcriptionally active genes and also involved in long-term transcriptional regulation in continuous white light (Benhamed et al., 2006; Guo et al., 2008; Gates et al., 2017).

We identified H3K9ac peaks in all samples, and differentially quantified them comparing wt D vs R1h, wt D vs wt WLc, and wt D vs *pifq* D. We found that the majority of the H3K9ac peaks mapped slightly downstream of the transcriptional start site region irrespective of light treatment, in concordance with previously published results for genome-wide H3K9ac profiles (Figure S4) (Charron et al., 2009). In dark-grown *pifq*, 7089 genes had statistically significantly higher H3K9ac levels than in wild type, while 900 genes had statistically lower H3K9ac levels (Supplemental Dataset S4). In WLc samples, 6797 genes had higher H3K9ac levels and 2505 genes had lower levels (Supplemental Dataset S4). These data are consistent with a change in the chromatin profile associated with photomorphogenic-like development in dark-grown *pifq* plants, where light-responsive genes become activated in the dark in the absence of the PIF quartet. In fact, clustering analysis revealed that the H3K9ac profile of *pifq* D is more similar to that of WLc-grown wild type than to dark-grown wild type or R1h-treated (Figure S5).

After one hour of red light, 992 genes had statistically significant higher H3K9ac levels than in dark-grown seedlings while 336 genes had lower H3K9ac (Supplemental Dataset S4). These results indicate that H3K9ac modification occurs only in a small fraction of light-regulated genes initially after light-exposure. H3K9ac levels change in a larger number of genes as light-exposure is sustained over longer periods, in a similar fashion to transcriptional changes. In every condition tested, H3K9ac levels generally correlated with mRNA levels (R^2^=0.46 for *pifq* vs D, R^2^=0.45 for R1h vs D and R^2^=0.64 for WLc vs D) for genes with a SSTF change in mRNA and H3K9ac levels (Figure S6).

To explore whether the PIF quartet are involved in the rapid red-light-induced chromatin changes, we focused our analysis of H3K9ac changes in LRP-DTGs in dark-grown *pifq* mutants, and in R1h-exposed wild type samples. We found that a high proportion of LRP-DTGs undergo H3K9ac changes that are associated with red-light induced PIF degradation (Figure 1B, 69% and 64% of PIF-induced LRP-DTGs and 68% and 63% of PIF-repressed LRP-DTGs in *pifq* and R1h respectively). In addition, the correlation between H3K9ac and mRNA levels was slightly higher when only LRP-DTGs were considered (R^2^=0.46 for *pifq* vs D, R^2^=0.65 for R1h vs D and R^2^=0.73 for WLc vs D, Figure S7). In summary, these results show that H3K9ac changes occur in the majority of LRP-DTGs, suggesting that it could be a key regulatory factor in their transcriptional regulation.

### H3K9 deacetylation parallels light-triggered transcriptional repression and apparently precedes transcriptional induction

Given the close parallel between the transcriptional and chromatin responses, we approached the question of whether this represents a causal relationship by performing concurrent time-course analysis of the light-induced responses in these two parameters. For this purpose, we selected six PIF-induced/light-repressed LRP-DTGs (*PIL1*, *XTR7*, *HB2*, *CKX5*, *IAA19* and *At5g02580*) and six PIF-repressed/light-induced LRP-DTGs (*RPT2*, *SIGE*, *LNK3*, *SPA1*, *BOH2* and *At1g60590*), based on the extent of their changes in mRNA and H3K9ac levels in the genome-wide experiments, and on their potential key roles in red-light responsiveness based on their molecular function (Figures 1C, 1D and S8, Table S1).

We first confirmed the results of the genome-wide analysis by measuring changes in H3K9ac levels in these 12 genes in response to R1h by ChIP-qPCR (Figure S9). We then performed a detailed time-course analysis of the rapid changes in mRNA and H3K9ac levels over the one-hour period following initial light exposure. We measured both parameters by RT-qPCR and ChIP-qPCR in the 12 selected LRP-DTGs, at 0, 1, 5, 10, 20, 30, 40, 50, and 60 minutes after a saturating red-light pulse. In broad terms, H3K9ac and mRNA levels were correlated (Figure S10). A more detailed analysis showed that the PIF-induced/light-repressed LRP-DTGs display rapid decreases in H3K9ac levels in response to red-light treatment, in parallel with the decrease in mRNA levels, starting just 5 minutes after the red-light pulse (Figures 2A and 2B). A converse pattern can be seen for the PIF-repressed/light-induced LRP-DTGs, where an increase in both mRNA and H3K9ac levels is detected shortly after red-light exposure (Figure 2C and 2D). However, in this case H3K9ac increase seems to occur earlier than mRNA induction, with detectable H3K9ac changes occurring around 10 minutes after the red-light pulse, while the mRNA increase happens 20 minutes after the re-light pulse (Figure 2C and D). These results suggest that H3K9ac deposition is necessary to initiate transcription, while its removal instantly triggers a reduction in steady state mRNA levels. It is also possible that this observed delay in transcription is caused by the fact that we are measuring steady-state levels of processed mRNA by RT-qPCR, and this technique does not capture the exact moment of transcriptional initiation.

**Figure 2.**
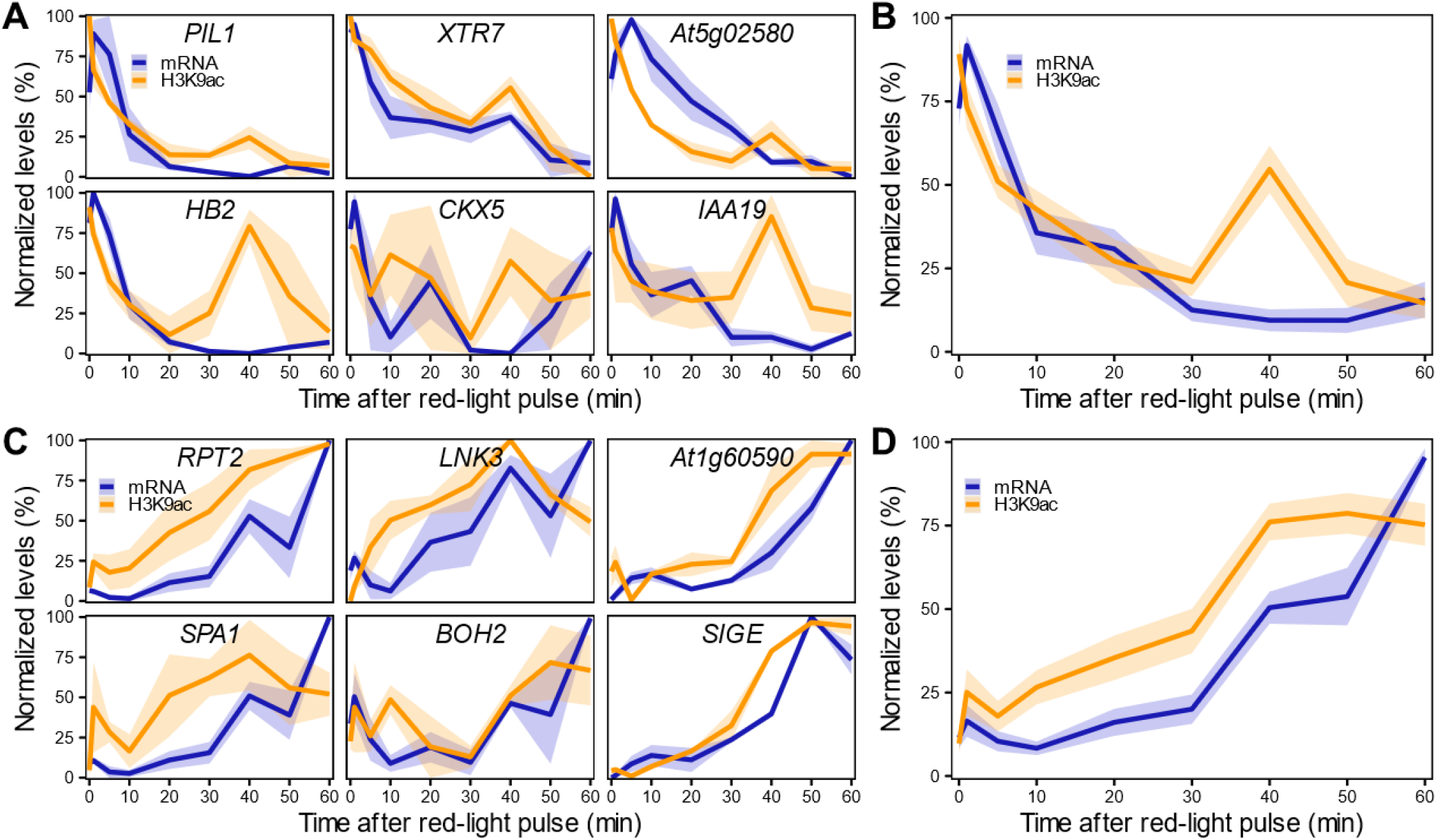
H3K9ac changes in LRP-DTGs parallel the rapid transcriptional response in light-repressed LRP-DTGs, and apparently precede transcription in light-induced LRP-DTGs. H3K9ac and mRNA changes measured by ChIP-qPCR and RT-qPCR in light-repressed (A) and light-induced (C) LRP-DTGs after a saturating red light pulse. Averaged mRNA and H3K9ac levels for each group of genes are shown in panels B and D, respectively. Data were re-scaled to the minimum and maximum mRNA/H3K9ac values for each gene. Each colored line represents the averaged mRNA/H3K9ac levels at each time point and the shaded band represents the standard error of the mean (n = 3).

### Real-time transcription imaging reveals an earlier timing of light-induced transcription initiation

In order to accurately measure a more continuous transcriptional readout and pinpoint the exact time of transcription initiation in response to light we generated several reporter lines. We first tested if we were able to recapitulate light-induced transcriptional initiation in transgenic lines expressing luciferase under the control of the *RPT2* promoter, a light-induced LRP-DTG (*pRPT2:LUC* lines). We chose *pRPT2* as it showed a strong and robust response to light in all our previous experiments. However, we were not able to detect any significant change in luminescence between *pRPT2:LUC* lines grown in the dark and after R1h treatment (Figure S11A). This absence of response was not due to *pRPT2* lacking light/PIF responsive elements, as we could detect a large luminescence increase when we transformed this reporter into *pifq* background (Figure S11B). It is likely that protein reporters that require transcription and translation in order to be measurable are not able to capture these short-term responses.

We then generated reporter lines in which we could measure mRNA transcription in real time (Figure S12). The *RPT2* promoter was used to drive transcription of a mRNA tagged in its 5’ with a PP7 sequence recognized by a co-expressed GFP-tagged PP7 phage coat protein, enabling identification of nascent mRNA as fluorescent puncta (*pRPT2:PP7* - Figure S12, (Larson et al., 2011; Alamos et al., 2021)). Imaging of dark-grown *pRPT2:PP7* lines exposed to light was able to recapitulate light-induced transcriptional initiation (Figure 3A). We measured real-time mRNA production in four independent *pRPT2:PP7* transgenic lines and we observed that transcription begins approximately 10 minutes after light exposure, slightly earlier than the increase in steady-state *RPT2* mRNA levels we detected by RT-qPCR (Figure 3B). The timing of transcription initiation coincides with the increase in H3K9ac levels observed by ChIP-qPCR (Figure 2C). Together, these results suggest that transcription initiation is accompanied by H3K9 acetylation, and, within the time resolution limitations of our experiments, could indicate that they are intertwined processes.

**Figure 3.**
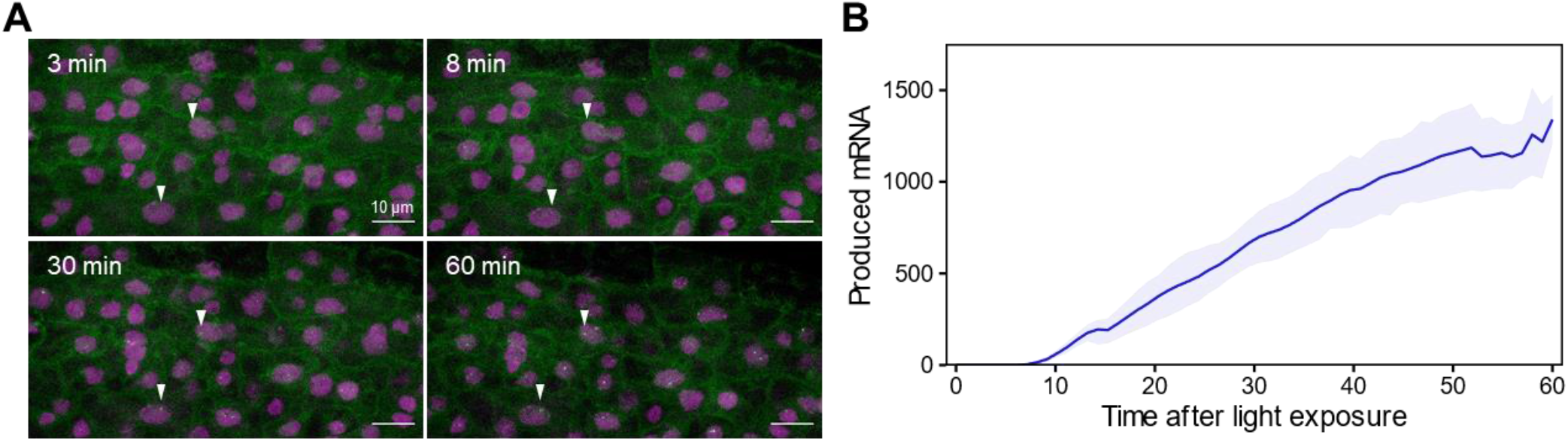
Single cell real-time transcription reveals an earlier start of light-induced *RPT2* transcription. A, Maximum projection snapshots of *pRPT2:PP7* plants grown in true-dark conditions for 2 days. The time stamp indicates the time elapsed since the seedlings were exposed to light. Arrowheads point to the appearance of transcription spots. B, Mean produced mRNA calculated as the integrated spot fluorescence over time. The colored line represents the average mRNA level and the shaded band represents the standard error of the mean. Data obtained from 2-5 technical replicates of four independent *pRPT2:PP7* transgenic lines are represented.

### H3K9ac writers and erasers involved in long-term light responses do not participate in fast H3K9ac changes in response to light

It has been previously reported that the transcriptional program of light-grown plants is disrupted in mutants for H3K9ac deposition (HAG1/GCN5 and HAF2/TAF1) and removal (HDA19/HD1) (Bertrand et al., 2005; Benhamed et al., 2006; Guo et al., 2008). To test the possibility that these H3K9ac writer or eraser proteins could be involved in gene regulation shortly after initial light-exposure, we measured mRNA levels of the 12 selected LRP-DTGs in dark-grown and R1h-treated *hag1*, *haf2* and *hda19* mutants. Contrary to what happens to other light-related genes in light-grown seedlings of these mutants, LRP-DTGs expression remained generally unaltered after a short red-light treatment. We only detected R1h-insensitive responses of a few LRP-DTGs (*At1g60590*, *LNK3* and *BOH2*), and only in the *hag1* mutants compared to wild type (Figure S12). These results indicate that these proteins, involved in H3K9ac regulation, and previously identified to have a role in long-term light-induced chromatin states and transcription are not involved in general short-term transcriptional regulation in response to light. Alternatively, another possible explanation is that because of functional redundancy of histone modifiers (Pandey et al., 2002), mutation in a single family member is not sufficient to affect the LRP-DTGs rapid transcriptional response to light. It remains to be explored which other possible factors might be responsible for the quick changes in H3K9ac in response to light exposure.

## Conclusions

The aim of the present study was to identify key genes that respond in the first instance to the first light exposure in plants, and understand the factors involved in their regulation. To do this, we focused on the red-light phytochrome signaling pathway. More specifically, centered on PIF direct target genes. We found that only a subset of the previously described PIF-DTGs respond most rapidly to light exposure. We redefined them as LRP-DTGs. These genes probably are the most direct targets of PIF regulation and play a key role in photomorphogenesis initiation. We found that the majority of these genes also show very rapid changes in H3K9ac levels, that accompany their transcriptional response, in the very initial minutes after light exposure. These findings suggest that chromatin remodeling is a crucial part of the very initial light response. This study contributes to our understanding of transcriptional regulation in response to environmental changes, and describes a system in which chromatin dynamics in response to environmental cues can be analyzed. The precise mechanism of how these histone modifications are performed remains to be elucidated.

## Materials & Methods

### Plant materials and growth conditions

The Columbia-0 ecotype of *Arabidopsis thaliana* was used for all experiments. *pif1pif3pif4pif5* (*pifq*) line is described in (Leivar et al., 2008), *hag1-5* (SALK_048427) in (Kornet and Scheres, 2009), *haf2-5* (SAIL_548_G03) in (Lee and Seo, 2018) and *hda19-4* (SALK_139443) in (Kim et al., 2008). To generate *pRPT2:LUC* lines, 3342 bp upstream of the RPT2 start codon were amplified by PCR with primers BamHI-pRPT2-5/ pRPT2-3-PstI and cloned into the pC1302-35S:RLUC vector (Zhang et al., 2013). To generate *pRPT2:PP7* lines, 3356 bp upstream of the RPT2 start codon were amplified by PCR with primers 13Rb-RPT2F and 13Rb-RPT2R and cloned into the AL13Rb plasmid (Addgene # 161006) (Alamos et al., 2021). Primers are described in Supplementary Dataset S7. Constructs were transformed into Arabidopsis by the floral dip method (Clough and Bent, 1998).

Plants were germinated in true-dark conditions as described in (Leivar et al., 2008). Briefly, seeds were sterilized, plated in MS medium without sucrose under white light and stratified for 4 days at 4°C in darkness. Afterwards, they were irradiated for 3h with white light to induce germination, followed by a saturating 5 min far-red light pulse. They were grown in the dark at 21°C for 2 days before being collected (D samples) or grown in the dark for 47 hours and treated with 10 μE·m^-2^·s^-1^ red light for 1 hour (R1h samples). White-light grown samples (WLc) were grown under continuous white-light (100 μE·m^-2^·s^-1^ PAR) after stratification for 2 days. For the time-course experiments, a saturating red-light pulse (5000 μE in 1 min) was given after 2 days in the dark and samples were collected at the indicated time-points.

### RNA sequencing

RNA-seq data from dark-grown wild-type and *pifq* mutants was obtained and analyzed previously by (Zhang et al., 2013).Total RNA from 2-day old “true dark”-grown (D), R1h and WLc treated seedlings was extracted and processed as described in (Zhang et al., 2013). RNA was extracted from 100 mg of ground tissue using RNeasy Plant Mini Kit (Qiagen cat. 74904) using RNase-Free DNase (Qiagen cat. 79254) following manufacturer instructions. The sequencing library construction was adapted from 3’-end RNA-seq protocol (Yoon and Brem, 2010) and performed as described in (Zhang et al., 2013). The size of purified library DNA was validated by Bioanalyzer 2000. Libraries from were assayed by 50-cycle single-end sequencing on the HiSeq2000 platform. Sequencing reads were aligned to the TAIR9 representative transcriptome using Bowtie (Langmead et al., 2009) with zero mismatches allowed. Only reads mapping uniquely to the 3’-end 500-bp region of the coding strand were counted for gene expression. Differentially expressed genes were identified using the edgeR package (Robinson et al., 2010), and SSTF genes were defined as those that differ by ≥2-fold with an adjusted P value ≤0.05. Sequencing data have been deposited in NCBI’s Gene Expression Omnibus (Edgar et al., 2002) and are accessible through GEO Series accession number GSE181167. Previously published sequencing data can be accessed through GEO Series accession number GSE39217.

### Chromatin Immunoprecipitation and sequencing

For each replicate, 2.5 gr of 2-day old wild-type and *pifq* D, R1h and WLc treated seedlings were processed as described in (Gendrel et al., 2005) using 5 μgr Diagenode Ab C15410004 H3K9ac antibody and a mix of 25/25 μl of protein-A/G dynabeads (Invitrogen). 5-10 ng of DNA per sample were used for library preparation with Accel-NGS 2S Plus DNA Library Kit (Swift Biosciences) and 9 PCR cycles. 300-700 bp fragments were purified and sequenced in an Illumina HiSeq 4000 by SR50 Single Read Sequencing. Reads were aligned to Arabidopsis thaliana TAIR10 genome using Bowtie2 (Langmead and Salzberg, 2012). H3K9ac enriched regions were identified using the BayesPeak algorithm with lower-bound summarization method (Spyrou et al., 2009). Differential enrichment analysis was performed with DiffBind (Stark and Brown, 2011), comparing H3K9ac enrichment in these regions among different samples. Only peaks with statistically significant different levels with an FDR ≤0.05 were used in the analysis. Sequencing data have been deposited in NCBI’s Gene Expression Omnibus (Edgar et al., 2002) and are accessible through GEO Series accession number GSE181432.

### Reverse transcription quantitative PCR

Total RNA was extracted from 100 mg of dark grown or R1h treated seedlings per biological replicate, and three biological replicates were used per time point using RNeasy Plant Mini Kit (Qiagen cat. 74904) with RNase-Free DNase (Qiagen cat. 79254) following manufacturer instructions. One μgr of RNA was used to make cDNA with the High-Capacity cDNA archive kit (Thermo cat. 4368814). qPCR reactions were performed with SYBR Green Master Mix (Thermo cat. 4309155) using three technical replicates per reaction in a CFX96 Touch Real-Time PCR Detection System (Biorad). *TUBULIN1* was used as a normalizer for ChIP-qPCR experiments and *PP2AA3* for RT-qPCR (Czechowski et al., 2005). Primers are described in Supplementary Dataset S7.

### Luciferase assays

Approximately 100 mg of two-day old seedlings were collected and ground in liquid nitrogen. Total protein was extracted using 100 μL of Passive Lysis Buffer (Promega). 20 μl of the supernatant were used to measure the LUC and RLUC activity using a Dual-Luciferase Reporter Assay System (Promega) according to the manufacturer’s instructions in a Tecan M-100PRO plate reader. Firefly luminescence was normalized by the constitutively expressed Renilla luciferase internal control.

### Real-time transcription imaging

Image acquisition and analysis was performed as described in (Alamos et al., 2021). Estimation of mRNA production was calculated as described in (Alamos et al., 2021) and (Garcia et al., 2013).

## Acknowledgements

We thank Robert H. Calderon and James Tepperman for fruitful discussions on data analysis. Figure 1A was created with BioRender.com. RNA- and ChIP-sequencing were performed by the Vincent J. Coates Genomics Sequencing Lab and the Functional Genomics Laboratory at the California Institute for Quantitative Biosciences at UC Berkeley.

## Author Contributions

E.G. and P.H.Q. conceived the project and wrote the manuscript. E.G., S.A., Y.Z., K.K.N, H.G.G. and P.H.Q, designed the experiments. E.G., Y.Z., S.A., J.D. performed the experiments. E.G. and S.A. performed the data analysis. All authors have edited and commented on the manuscript and have given their approval of the final version.

