## Supplemental Figures for "Chromatin changes in PIF-regulated genes parallel their rapid transcriptional response to light"

### Slide 1
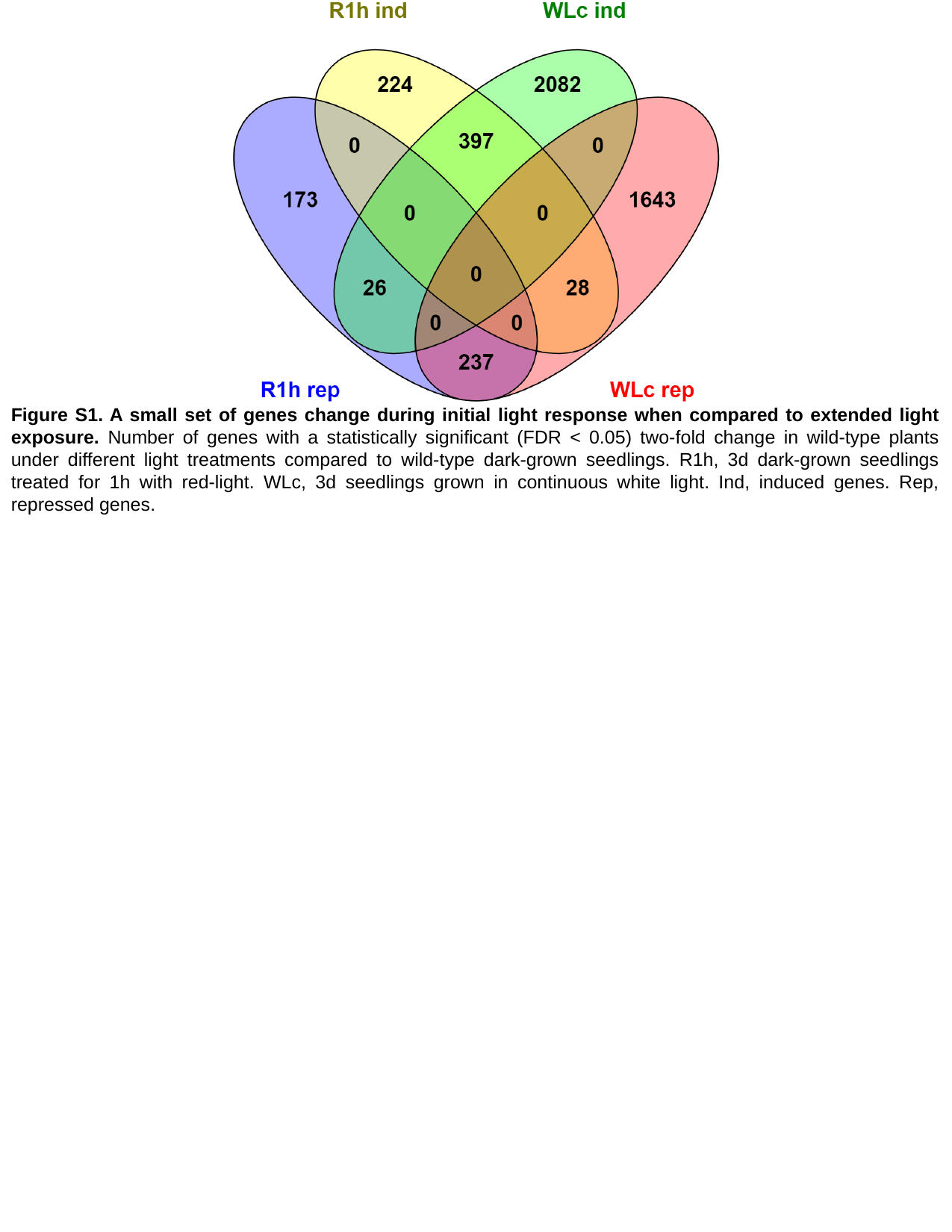

Figure S1. A small set of genes change during initial light response when compared to extended light exposure. Number of genes with a statistically significant (FDR < 0.05) two-fold change in wild-type plants under different light treatments compared to wild-type dark-grown seedlings. R1h, 3d dark-grown seedlings treated for 1h with red-light. WLc, 3d seedlings grown in continuous white light. Ind, induced genes. Rep, repressed genes.

### Slide 2
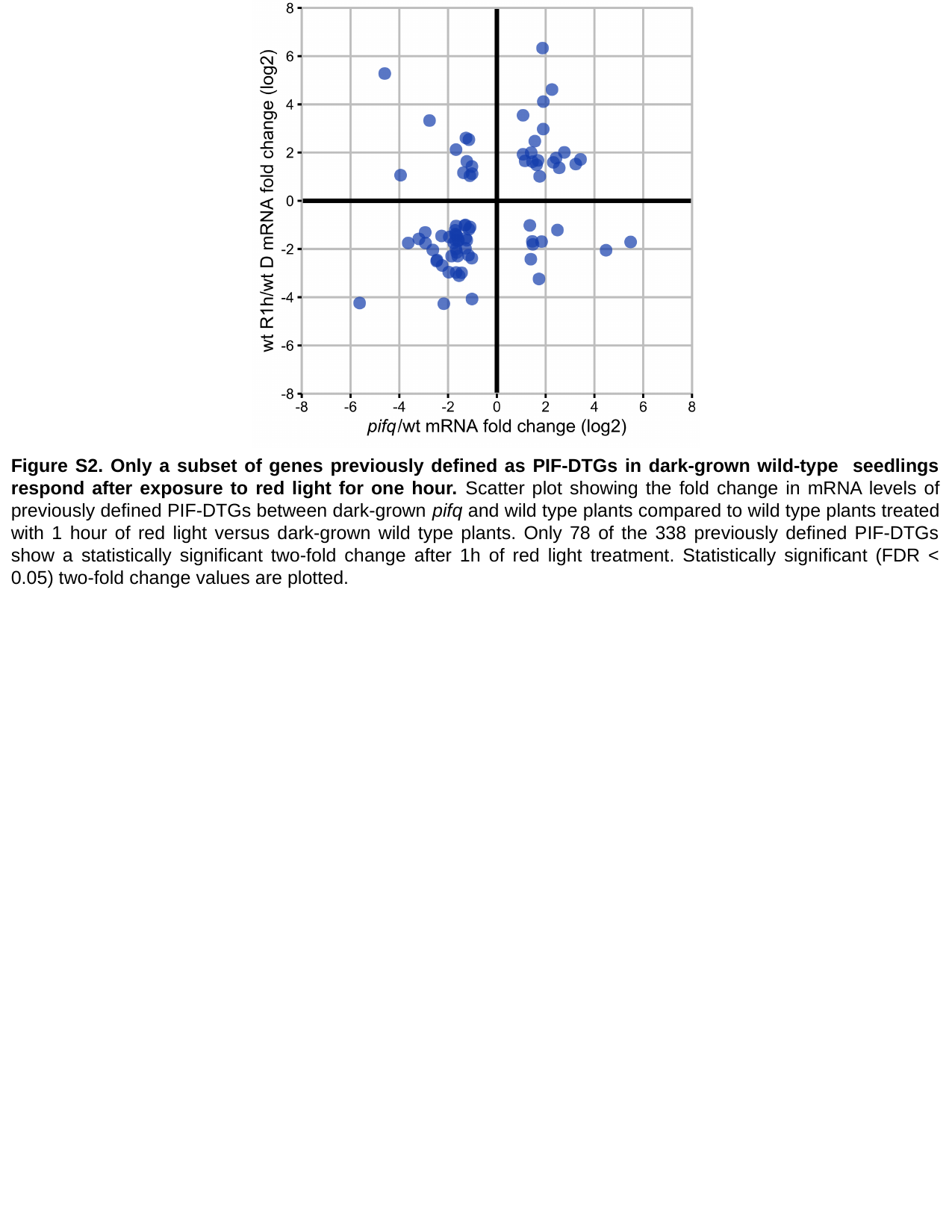

Figure S2. Only a subset of genes previously defined as PIF-DTGs in dark-grown wild-type seedlings respond after exposure to red light for one hour. Scatter plot showing the fold change in mRNA levels of previously defined PIF-DTGs between dark-grown pifq and wild type plants compared to wild type plants treated with 1 hour of red light versus dark-grown wild type plants. Only 78 of the 338 previously defined PIF-DTGs show a statistically significant two-fold change after 1h of red light treatment. Statistically significant (FDR < 0.05) two-fold change values are plotted.

### Slide 3
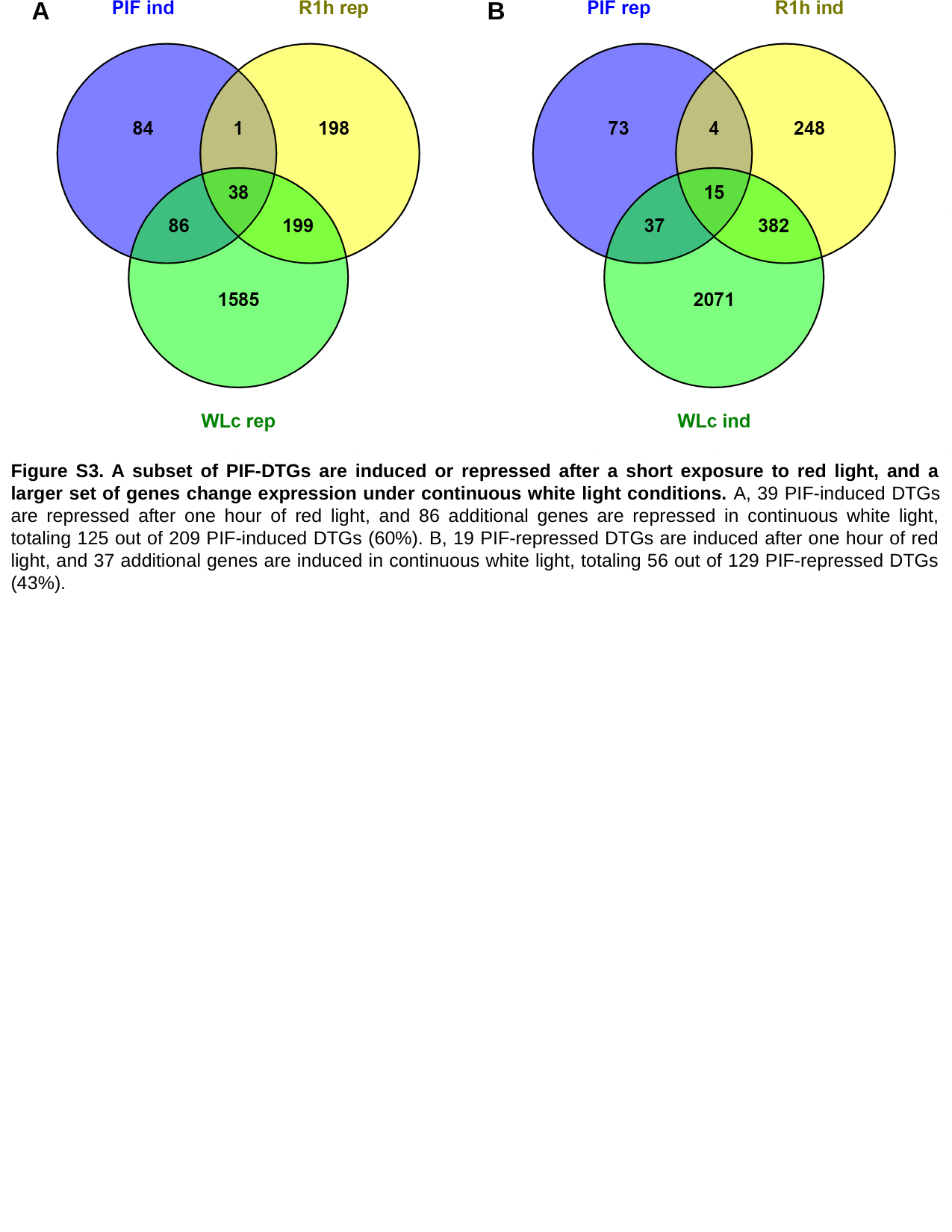

A
B
Figure S3. A subset of PIF-DTGs are induced or repressed after a short exposure to red light, and a larger set of genes change expression under continuous white light conditions. A, 39 PIF-induced DTGs are repressed after one hour of red light, and 86 additional genes are repressed in continuous white light, totaling 125 out of 209 PIF-induced DTGs (60%). B, 19 PIF-repressed DTGs are induced after one hour of red light, and 37 additional genes are induced in continuous white light, totaling 56 out of 129 PIF-repressed DTGs (43%).

### Slide 4
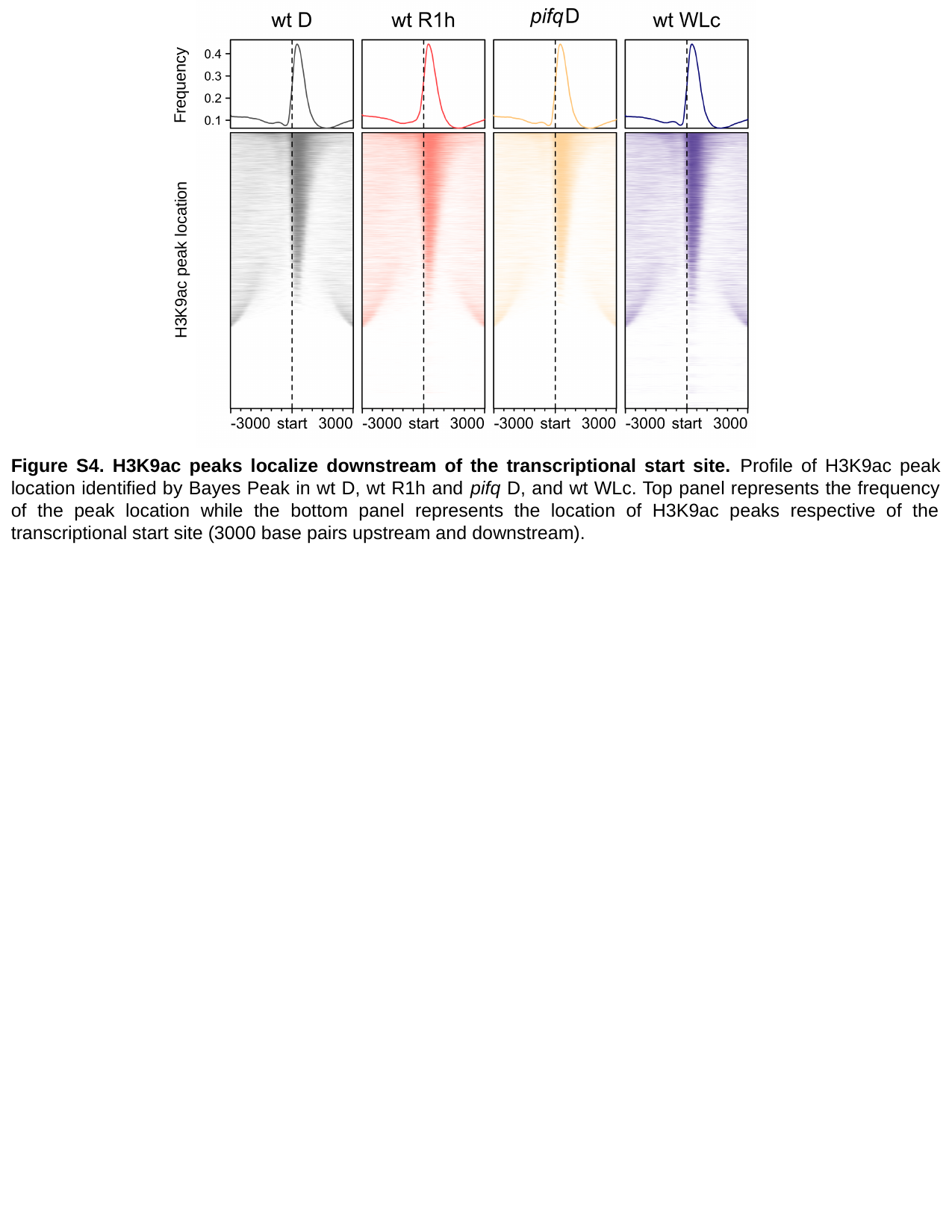

Frequency
H3K9ac peak location
Figure S4. H3K9ac peaks localize downstream of the transcriptional start site. Profile of H3K9ac peak location identified by Bayes Peak in wt D, wt R1h and pifq D, and wt WLc. Top panel represents the frequency of the peak location while the bottom panel represents the location of H3K9ac peaks respective of the transcriptional start site (3000 base pairs upstream and downstream).

### Slide 5
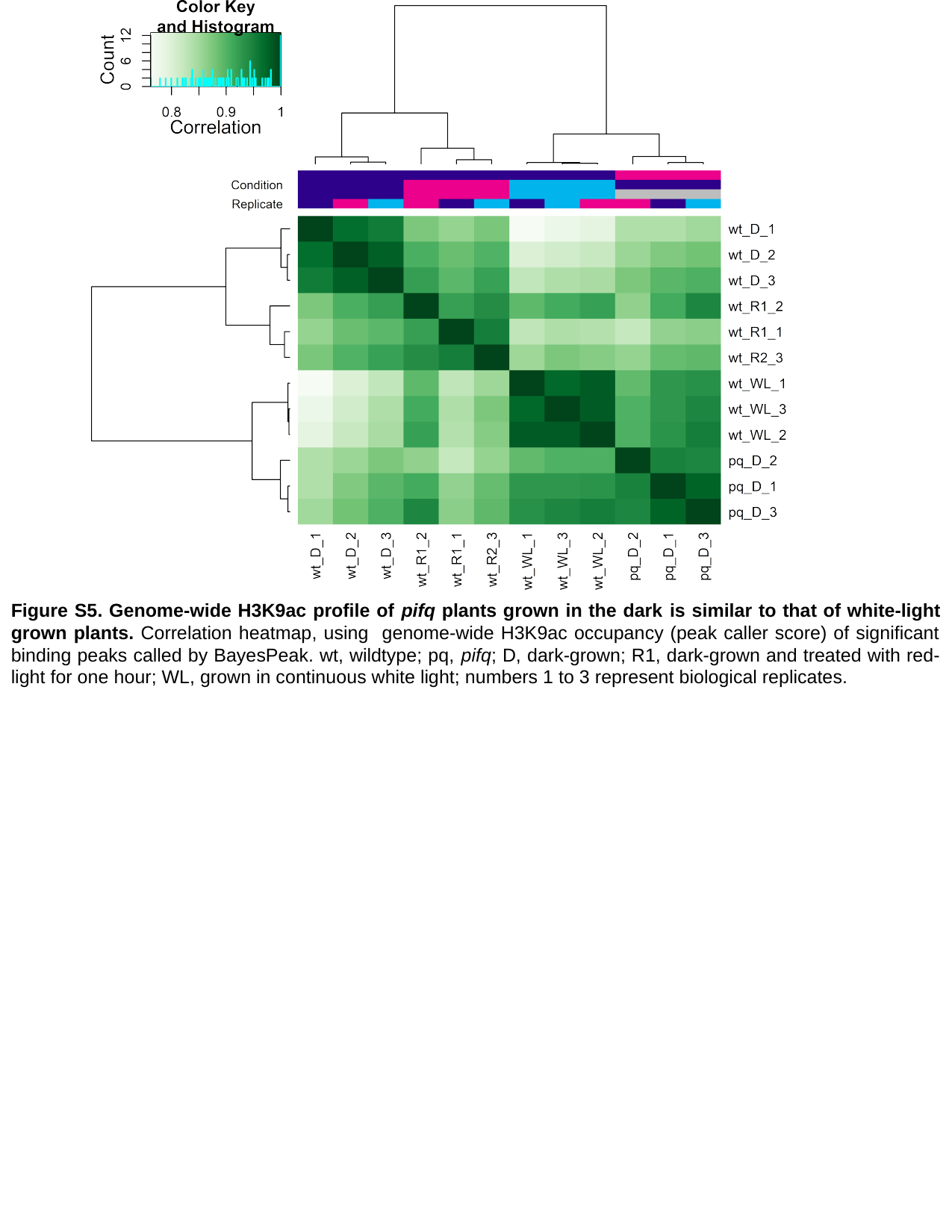

Figure S5. Genome-wide H3K9ac profile of pifq plants grown in the dark is similar to that of white-light grown plants. Correlation heatmap, using genome-wide H3K9ac occupancy (peak caller score) of significant binding peaks called by BayesPeak. wt, wildtype; pq, pifq; D, dark-grown; R1, dark-grown and treated with red-light for one hour; WL, grown in continuous white light; numbers 1 to 3 represent biological replicates.

### Slide 6
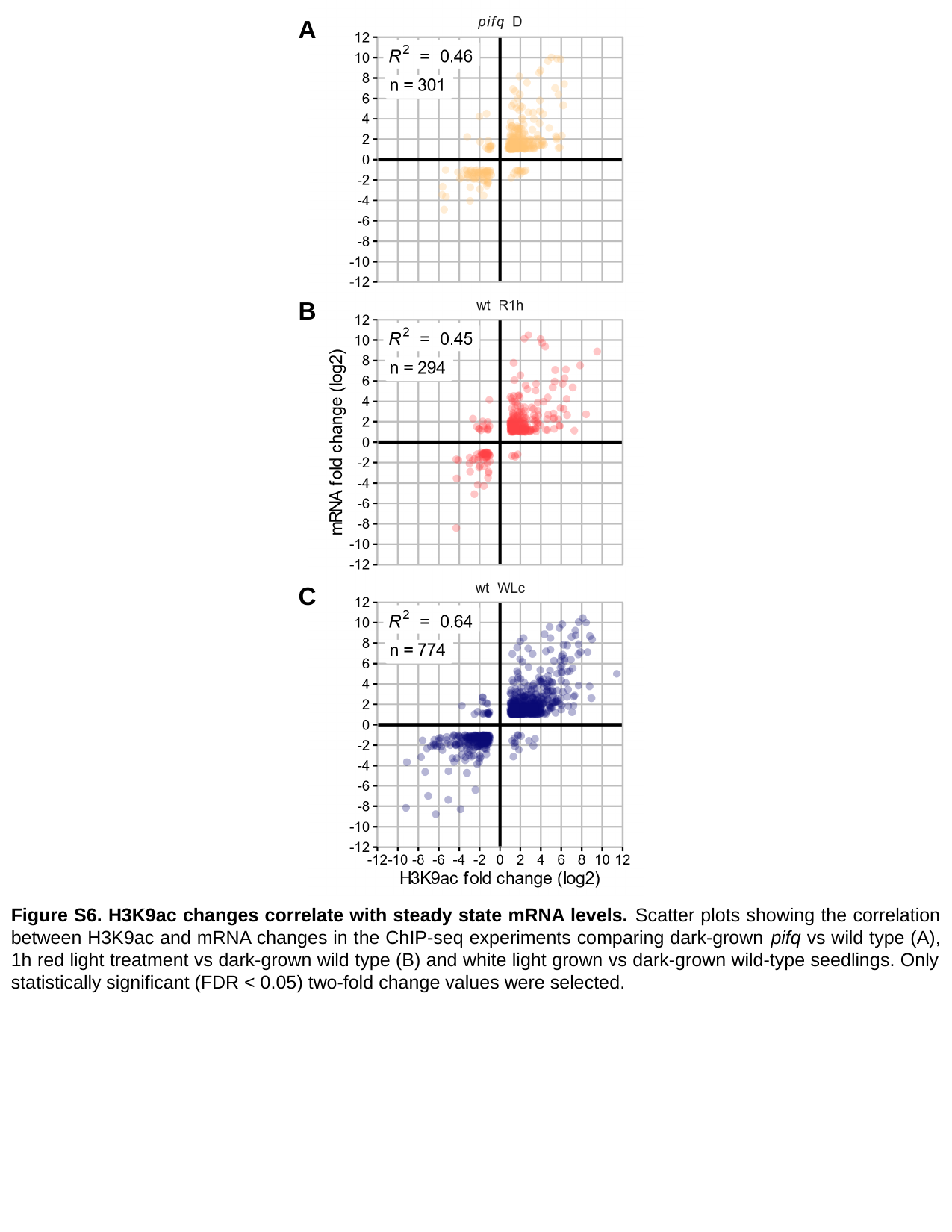

A
B
C
Figure S6. H3K9ac changes correlate with steady state mRNA levels. Scatter plots showing the correlation between H3K9ac and mRNA changes in the ChIP-seq experiments comparing dark-grown pifq vs wild type (A), 1h red light treatment vs dark-grown wild type (B) and white light grown vs dark-grown wild-type seedlings. Only statistically significant (FDR < 0.05) two-fold change values were selected.

### Slide 7
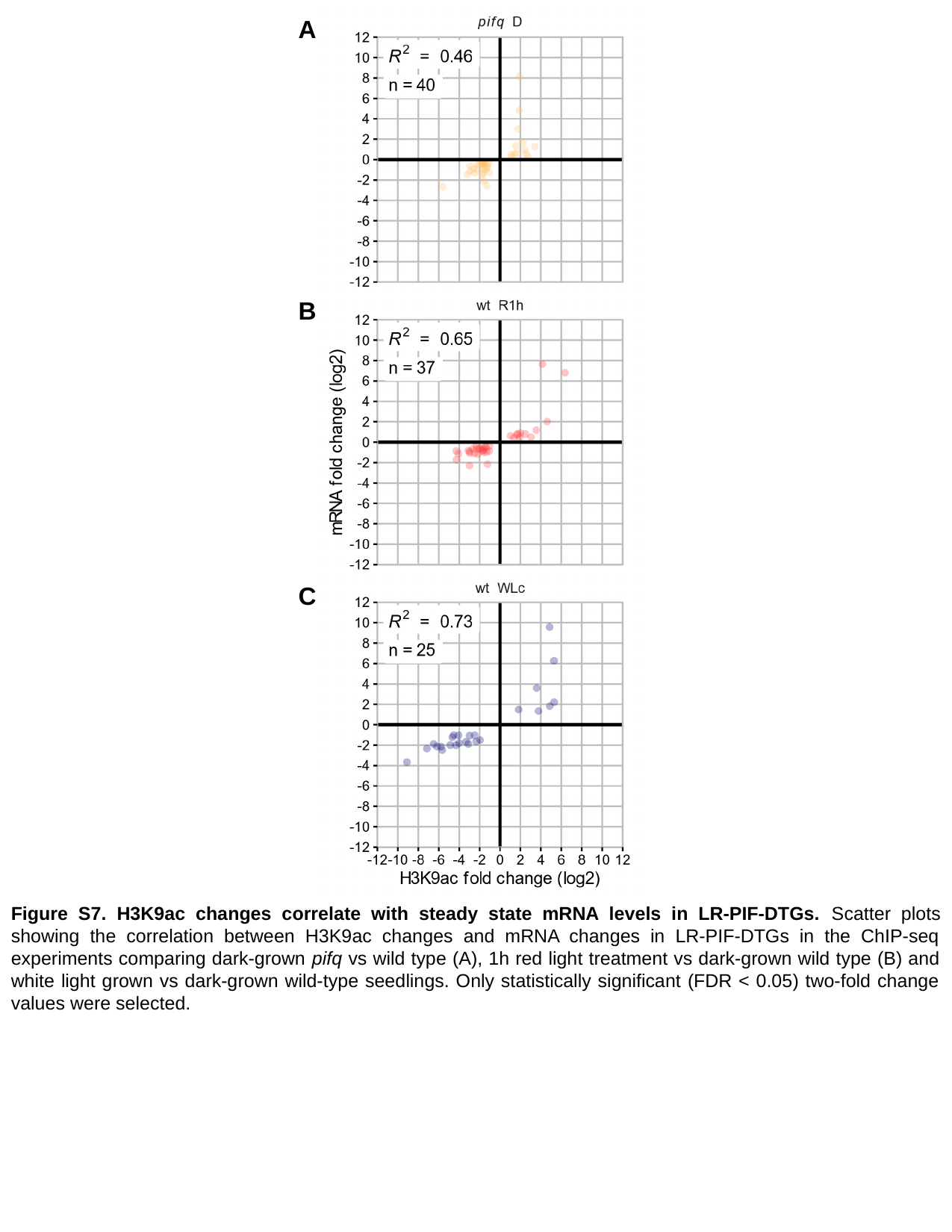

A
B
C
Figure S7. H3K9ac changes correlate with steady state mRNA levels in LR-PIF-DTGs. Scatter plots showing the correlation between H3K9ac changes and mRNA changes in LR-PIF-DTGs in the ChIP-seq experiments comparing dark-grown pifq vs wild type (A), 1h red light treatment vs dark-grown wild type (B) and white light grown vs dark-grown wild-type seedlings. Only statistically significant (FDR < 0.05) two-fold change values were selected.

### Slide 8
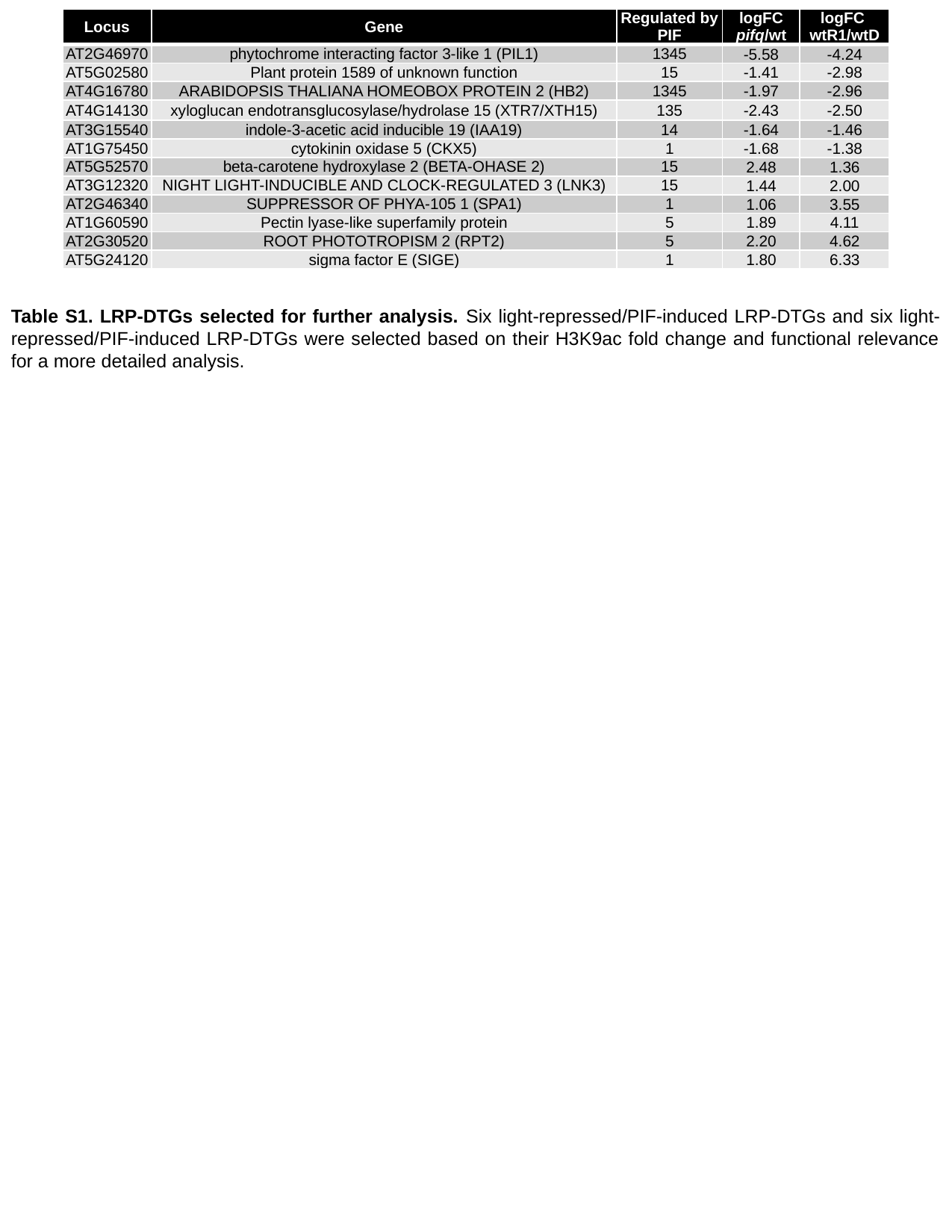

| Locus | Gene | Regulated by PIF | logFC pifq/wt | logFC wtR1/wtD |
| --- | --- | --- | --- | --- |
| AT2G46970 | phytochrome interacting factor 3-like 1 (PIL1) | 1345 | -5.58 | -4.24 |
| AT5G02580 | Plant protein 1589 of unknown function | 15 | -1.41 | -2.98 |
| AT4G16780 | ARABIDOPSIS THALIANA HOMEOBOX PROTEIN 2 (HB2) | 1345 | -1.97 | -2.96 |
| AT4G14130 | xyloglucan endotransglucosylase/hydrolase 15 (XTR7/XTH15) | 135 | -2.43 | -2.50 |
| AT3G15540 | indole-3-acetic acid inducible 19 (IAA19) | 14 | -1.64 | -1.46 |
| AT1G75450 | cytokinin oxidase 5 (CKX5) | 1 | -1.68 | -1.38 |
| AT5G52570 | beta-carotene hydroxylase 2 (BETA-OHASE 2) | 15 | 2.48 | 1.36 |
| AT3G12320 | NIGHT LIGHT-INDUCIBLE AND CLOCK-REGULATED 3 (LNK3) | 15 | 1.44 | 2.00 |
| AT2G46340 | SUPPRESSOR OF PHYA-105 1 (SPA1) | 1 | 1.06 | 3.55 |
| AT1G60590 | Pectin lyase-like superfamily protein | 5 | 1.89 | 4.11 |
| AT2G30520 | ROOT PHOTOTROPISM 2 (RPT2) | 5 | 2.20 | 4.62 |
| AT5G24120 | sigma factor E (SIGE) | 1 | 1.80 | 6.33 |
Table S1. LRP-DTGs selected for further analysis. Six light-repressed/PIF-induced LRP-DTGs and six light-repressed/PIF-induced LRP-DTGs were selected based on their H3K9ac fold change and functional relevance for a more detailed analysis.

### Slide 9
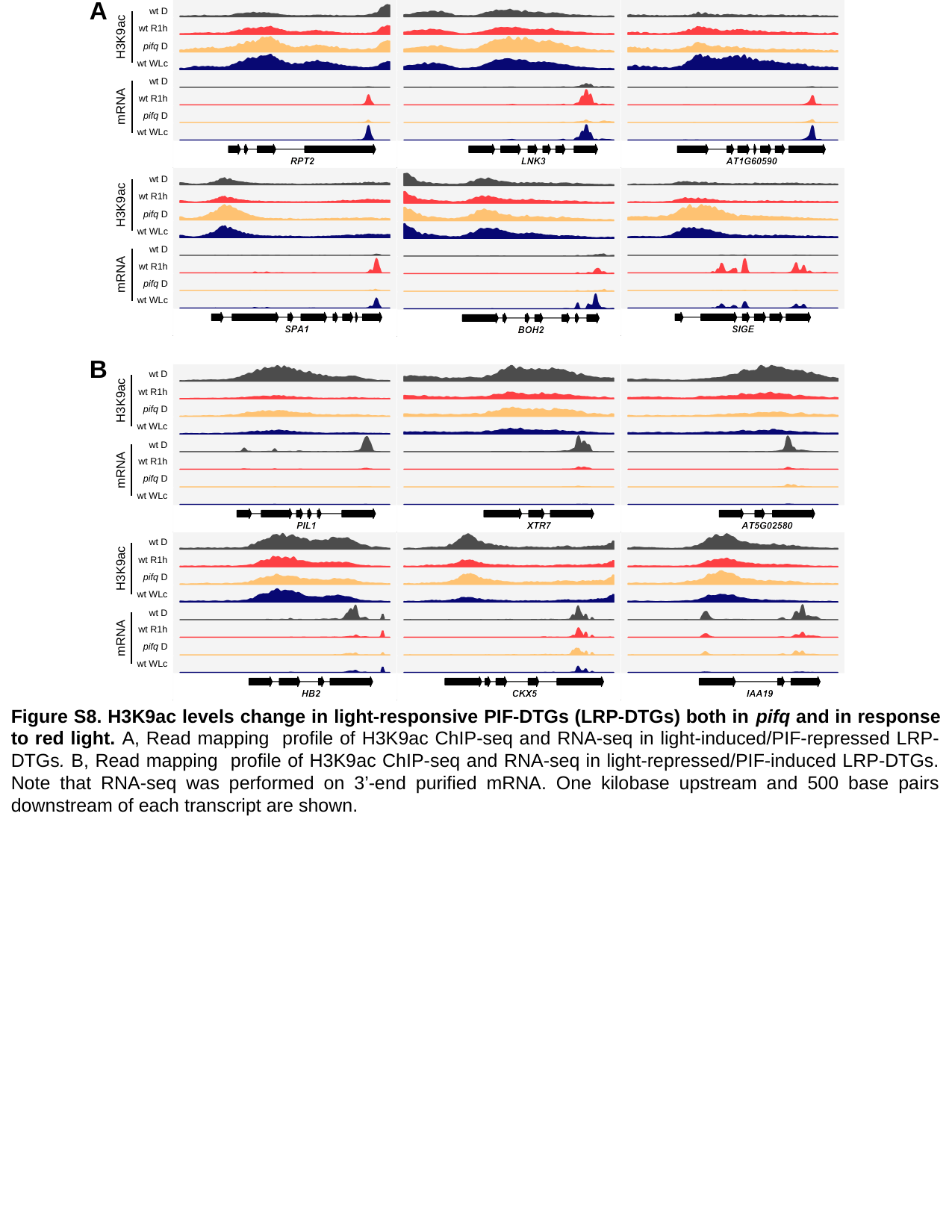

A
wt D
H3K9ac
wt R1h
pifq D
wt WLc
wt D
mRNA
wt R1h
pifq D
wt WLc
wt D
H3K9ac
wt R1h
pifq D
wt WLc
wt D
mRNA
wt R1h
pifq D
wt WLc
B
wt D
H3K9ac
wt R1h
pifq D
wt WLc
wt D
mRNA
wt R1h
pifq D
wt WLc
wt D
H3K9ac
wt R1h
pifq D
wt WLc
wt D
mRNA
wt R1h
pifq D
wt WLc
Figure S8. H3K9ac levels change in light-responsive PIF-DTGs (LRP-DTGs) both in pifq and in response to red light. A, Read mapping profile of H3K9ac ChIP-seq and RNA-seq in light-induced/PIF-repressed LRP-DTGs. B, Read mapping profile of H3K9ac ChIP-seq and RNA-seq in light-repressed/PIF-induced LRP-DTGs. Note that RNA-seq was performed on 3’-end purified mRNA. One kilobase upstream and 500 base pairs downstream of each transcript are shown.

### Slide 10
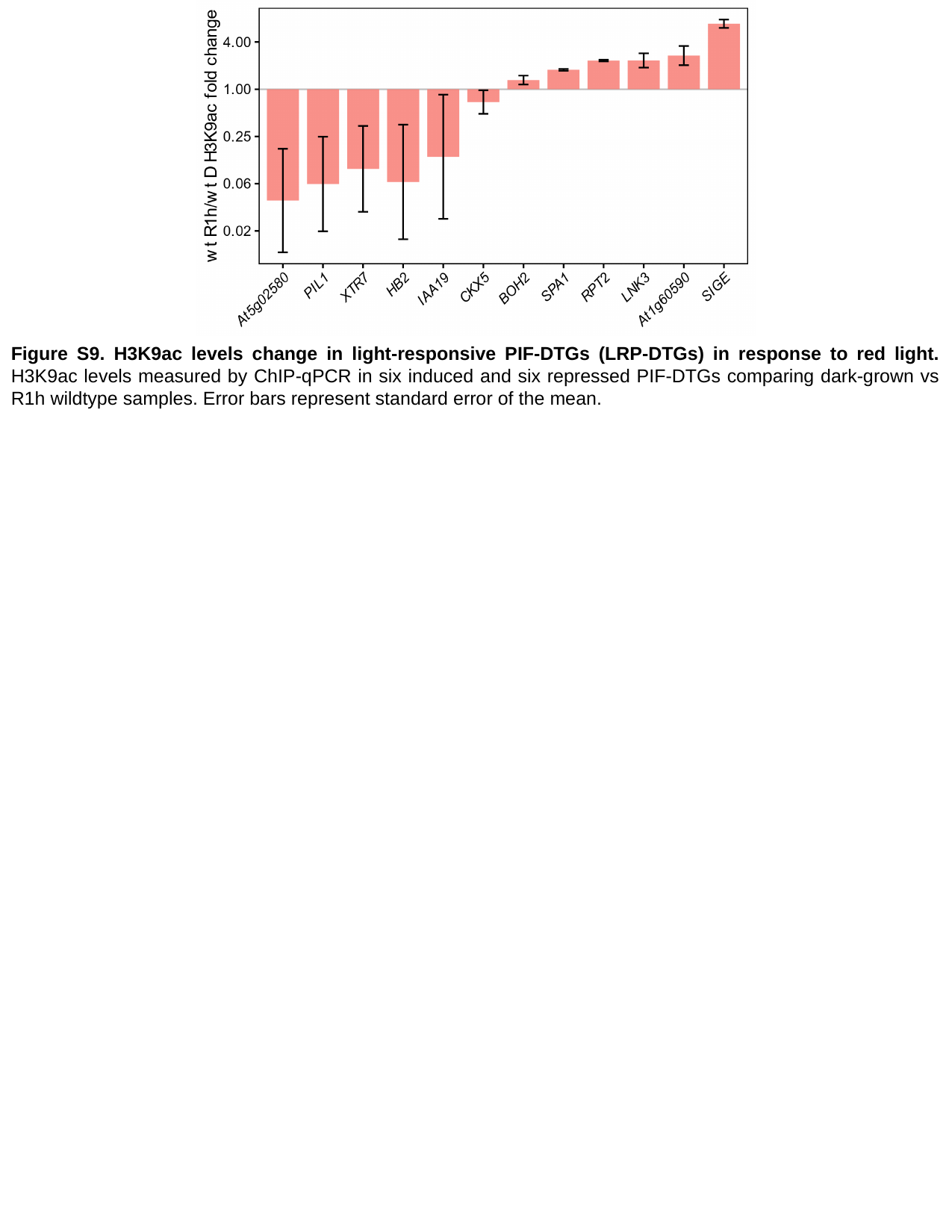

Figure S9. H3K9ac levels change in light-responsive PIF-DTGs (LRP-DTGs) in response to red light. H3K9ac levels measured by ChIP-qPCR in six induced and six repressed PIF-DTGs comparing dark-grown vs R1h wildtype samples. Error bars represent standard error of the mean.

### Slide 11
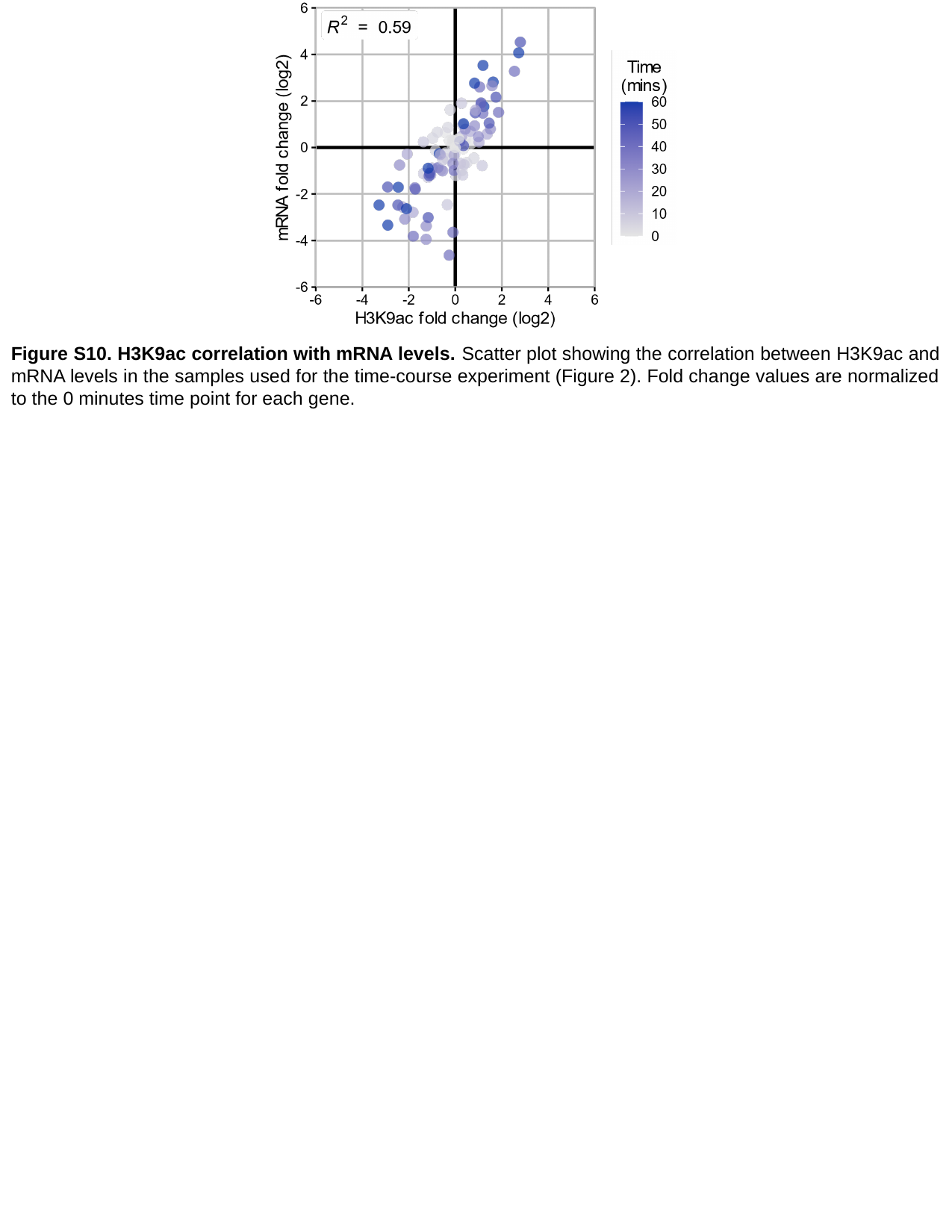

Figure S10. H3K9ac correlation with mRNA levels. Scatter plot showing the correlation between H3K9ac and mRNA levels in the samples used for the time-course experiment (Figure 2). Fold change values are normalized to the 0 minutes time point for each gene.

### Slide 12
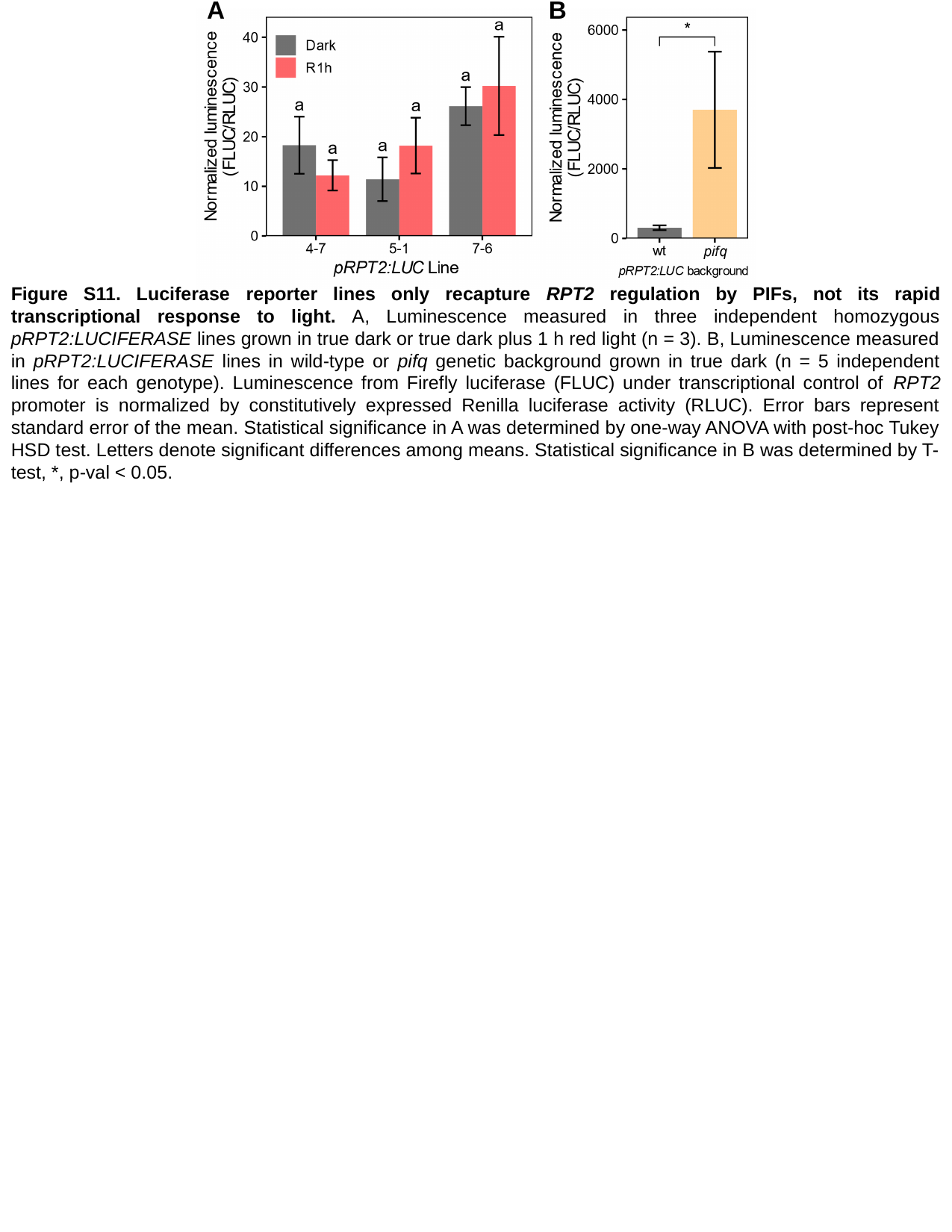

A
B
Figure S11. Luciferase reporter lines only recapture RPT2 regulation by PIFs, not its rapid transcriptional response to light. A, Luminescence measured in three independent homozygous pRPT2:LUCIFERASE lines grown in true dark or true dark plus 1 h red light (n = 3). B, Luminescence measured in pRPT2:LUCIFERASE lines in wild-type or pifq genetic background grown in true dark (n = 5 independent lines for each genotype). Luminescence from Firefly luciferase (FLUC) under transcriptional control of RPT2 promoter is normalized by constitutively expressed Renilla luciferase activity (RLUC). Error bars represent standard error of the mean. Statistical significance in A was determined by one-way ANOVA with post-hoc Tukey HSD test. Letters denote significant differences among means. Statistical significance in B was determined by T-test, *, p-val < 0.05.

### Slide 13
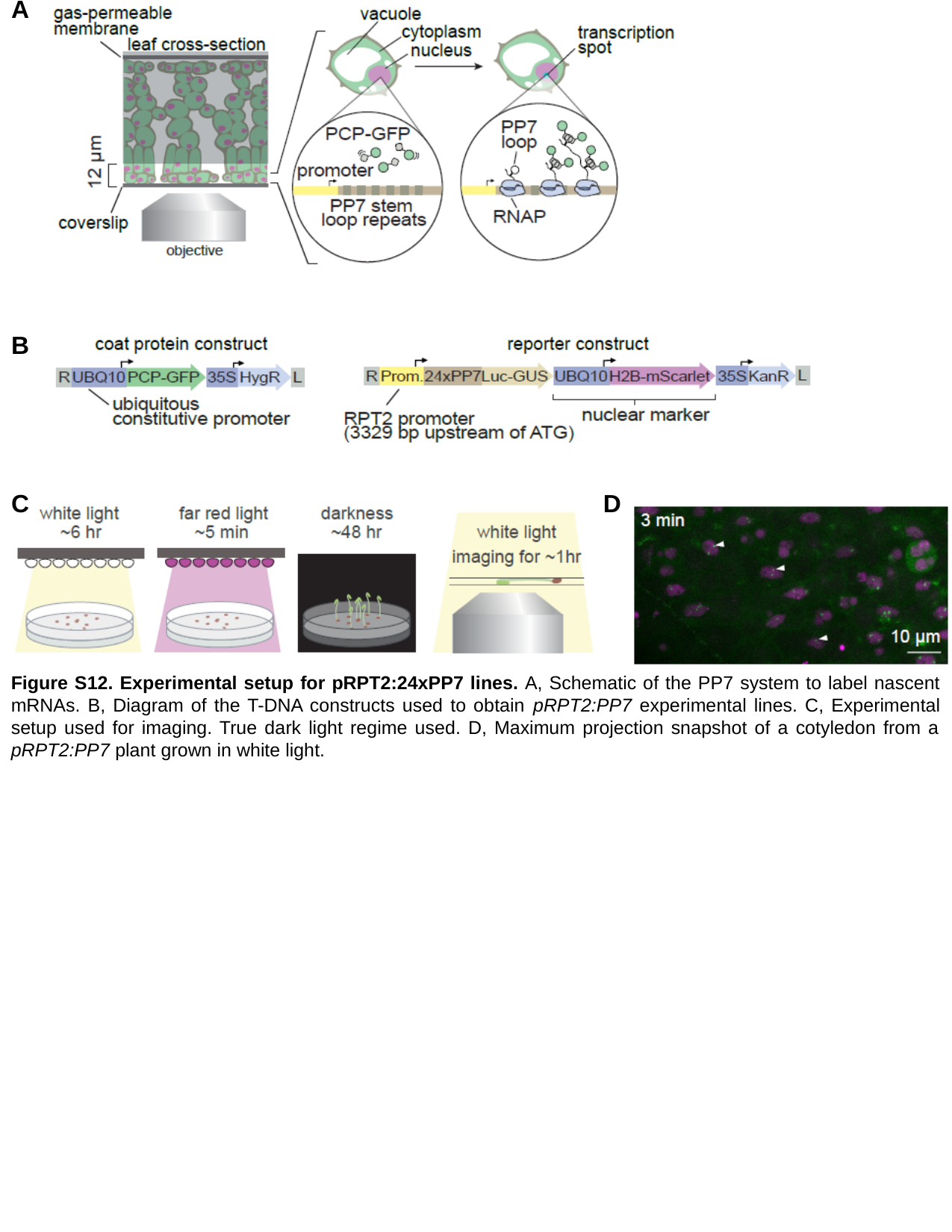

A
B
C
D
Figure S12. Experimental setup for pRPT2:24xPP7 lines. A, Schematic of the PP7 system to label nascent mRNAs. B, Diagram of the T-DNA constructs used to obtain pRPT2:PP7 experimental lines. C, Experimental setup used for imaging. True dark light regime used. D, Maximum projection snapshot of a cotyledon from a pRPT2:PP7 plant grown in white light.

### Slide 14
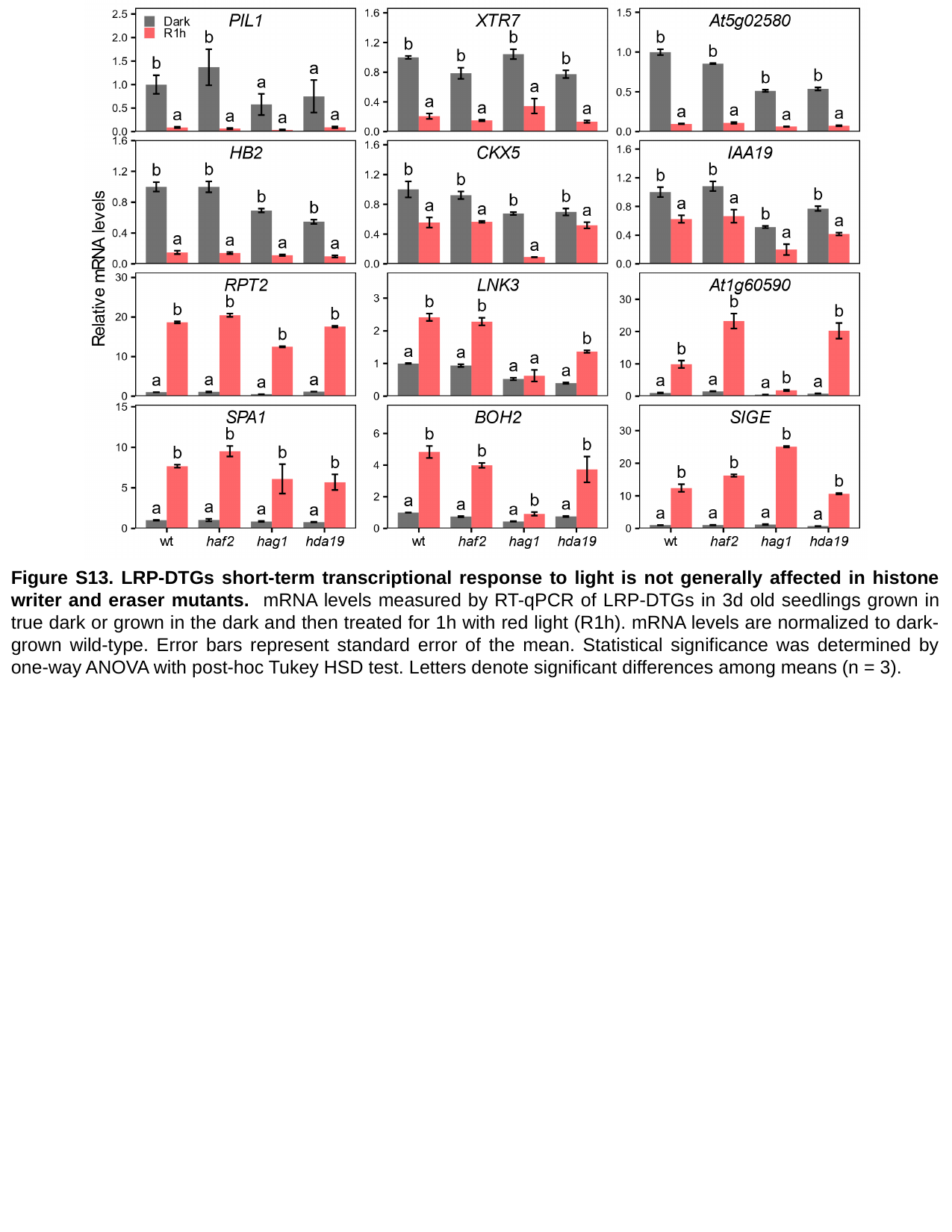

Figure S13. LRP-DTGs short-term transcriptional response to light is not generally affected in histone writer and eraser mutants. mRNA levels measured by RT-qPCR of LRP-DTGs in 3d old seedlings grown in true dark or grown in the dark and then treated for 1h with red light (R1h). mRNA levels are normalized to dark-grown wild-type. Error bars represent standard error of the mean. Statistical significance was determined by one-way ANOVA with post-hoc Tukey HSD test. Letters denote significant differences among means (n = 3).
